# N-1-naphthylphthalamic acid stimulates tomato hypocotyl elongation, elevates ethylene levels and alters metabolic homeostasis

**DOI:** 10.1101/311159

**Authors:** Sapana Nongmaithem, Sameera Devulapalli, Yellamaraju Sreelakshmi, Rameshwar Sharma

**Author notes:** **Corresponding author:** Prof. Rameshwar Sharma, Repository of Tomato Genomics Resources, Department of Plant University of Hyderabad, Hyderabad-500046, India.

## Abstract

**One sentence summary:** N-1-naphthylphthalamic acid (NPA) treatment stimulates tomato hypocotyl elongation likely by elevating ethylene emission and lowering indole-3-butyric acid levels in the seedlings.

**Abstract:** In higher plants, phytohormone indole-3-acetic acid is characteristically transported from the apex towards the base of the plant, termed as polar auxin transport (PAT). Among the inhibitors blocking PAT, N-1-naphthylphthalamic acid (NPA) that targets ABCB transporters is most commonly used. NPA-treated light-grown Arabidopsis seedlings show severe inhibition of hypocotyl and root elongation. In light-grown tomato seedlings, NPA inhibited root growth, but contrary to Arabidopsis stimulated hypocotyl elongation. The NPA-stimulation of hypocotyl elongation was milder in blue, red, and far-red light-grown seedlings. The NPA-treatment stimulated emission of ethylene from the seedlings. The scrubbing of ethylene by mercuric perchlorate reduced NPA-stimulated hypocotyl elongation. NPA action on hypocotyl elongation was antagonized by 1-methylcyclopropene, an inhibitor of ethylene action. NPA-treated seedlings had reduced levels of indole-3-butyric acid and higher levels of zeatin in the shoots. NPA did not alter indole-3-acetic levels in shoots. The analysis of metabolic networks indicated that NPA-treatment induced moderate shifts in the networks compared to exogenous ethylene that induced a drastic shift in metabolic networks. Our results indicate that in addition to ethylene, NPA-stimulated hypocotyl elongation in tomato may also involve zeatin and indole-3-butyric acid. Our results indicate that NPA-mediated physiological responses may vary in a species-specific fashion.

## Introduction

Plant development is mediated by coordinated synthesis and distribution of several plant hormones. Among the plant hormones, the auxin is involved in almost all developmental responses throughout the lifecycle of the plants (Zažímalová et al., 2014). While most plant hormones function in a cell-autonomous fashion, several developmental responses mediated by the auxins such as leaf primordia initiation, tropic movement of organs are attributed to its directional transport within a tissue and/or organ. The orientation of auxin transport in an organ or tissue is mediated by a battery of transporters and associated proteins, which together imparts the specialized transport termed as polar auxin transport (PAT) (Chen and Baluška, 2013). It is believed that auxin being a weak acid diffuses in the cells as an uncharged molecule, wherein it is dissociated to the charged form, and its efflux is mediated from the cells by polarly localized auxin transporters.

The analysis of the mutants defective in PAT led to the identification of several genes encoding for the proteins facilitating PAT. Several physiological and molecular evidences indicate that the PAT is mediated by the distinct influx and efflux carriers presumably localized at the plasma membrane. Molecular-genetic evidence indicated that the members of the AUX/LAX family act as auxin influx carriers (Péret et al., 2012). The auxin efflux from cells is mediated by a group of proteins belonging to PIN and ABCB family; and is modulated by additional proteins such as PINOID (Armengot et al., 2016). The maintenance of polar auxin transport system is essential for normal development of the plant. Thus, the mutants altered in auxin transport display several developmental abnormalities (Al-Hammadi et al., 2003; Okada et al., 1991).

Prior to molecular genetic analysis of mutants, most information regarding the contribution of auxin to plant development came from the use of pharmacological inhibitors that specifically affect auxin transport. The inhibitors like 1-naphthoxyacetic acid, 2-naphthoxyacetic acid, 3-chloro-4-hydroxyphenylacetic acid inhibit auxin influx carriers (Imhoff et al., 2000), whereas 1-N-naphthylphthalamic acid (NPA), 2-(1-pyrenoyl)-benzoic acid, cyclopropyl propane dione, 2,3,5-triiodobenzoic acid, morphactins inhibit auxin efflux carriers (Klíma et al., 2016). Among these inhibitors, NPA is the most widely used and classified as a phytotropin, a class of chemicals that inhibit tropic responses in stems. Using NPA, several developmental responses have been attributed to the PAT such as leaf vein patterning (Mattsson et al., 1999), phyllotaxy (Reinhardt et al., 2000), lateral root development (Casimiro et al., 2001) and embryo development (Schiavone and Cooke, 1987). The action of NPA on auxin efflux is attributed to its interaction with several cellular components. It binds to twisted-dwarf-1 (TWD1), ABCB1, ABCB19 and its binding to TWD1 and ABCB1 prevents their protein-protein interaction leading to reduced PAT (Noh et al., 2001). In addition, it also binds to plasma membrane-associated aminopeptidases APM1 and APP1 (Murphy et al., 2002). While NPA does not bind to PIN proteins, it alters the intracellular cycling of PIN proteins between endosomal vesicles and the plasma membrane (Dhonukshe et al., 2008; Geisler et al., 2005).

The auxin-induced cell expansion is one of the most extensively studied growth response. Since post-germination Arabidopsis hypocotyl growth mainly consists of cell expansion, several studies used hypocotyls as a model system to examine the role of light and phytohormones, particularly auxin and ethylene on hypocotyl elongation (Vandenbussche et al., 2005; Hu et al., 2017). Ethylene suppresses elongation of hypocotyls in dark, while it promotes elongation in light-grown seedlings (Smalle et al., 1997; Zhong et al., 2012). In light, ethylene stimulates translocation of COP1 to the nucleus where it mediates HY5 degradation contributing to hypocotyl growth in the light (Yu et al., 2013). The auxin transport plays an important role in hypocotyl elongation of light-grown seedlings. NPA treatment strongly inhibited hypocotyl elongation in light-grown seedlings whereas it had minimal effect in etiolated Arabidopsis seedlings (Jensen et al., 1998). The above light effect is likely related to auxin transport, as the rate of PAT is lower in hypocotyls of etiolated seedlings of Arabidopsis and tomato than in light-grown seedlings (Rashotte et al., 2003, Liu et al., 2012). The light-dependent auxin transport is likely mediated by both phytochrome and cryptochrome as NPA effect on hypocotyl elongation is subdued in the mutants defective in above photoreceptors (Jansen et al., 1998).

Relatively little information is available about the effect of NPA on the cellular metabolism and other plant hormones. Emerging evidence indicates that NPA also influences other hormonal responses. Arabidopsis plants grown on NPA containing media for five days showed significantly reduced indole-3-acetic acid (IAA) levels in the leaves (Ljung et al., 2001). A likely link between auxin transport and cytokinin level is indicated as NPA strongly decreased transcripts of cytokinin oxidases, *AtCKX1* and *AtCKX6* (Werner et al., 2006). Application of NPA mimicked the cytokinin induction of the off-the-medium growth of Arabidopsis root tip (Kushwah et al., 2011). NPA also mimicked the effect of exogenous cytokinin in inducing root like organogenesis in excised hypocotyls of Arabidopsis (Pernisová et al., 2009). A relationship between NPA and ethylene is also indicated as NPA sensitized roots of ethylene resistant *eir1* mutant to ethylene (Růžička et al., 2007).

To better understand the effect of NPA on plant hormones level and the cellular metabolism, we grew tomato seedlings on the medium containing NPA. Surprisingly, NPA stimulated hypocotyl elongation in tomato seedlings rather than inhibiting it as reported in Arabidopsis. This variance in NPA response compared to Arabidopsis was restricted to hypocotyl, as NPA inhibited root elongation akin to Arabidopsis. NPA treatment also influenced the metabolic and hormonal homeostasis of tomato seedlings, elevating ethylene synthesis, zeatin levels and reducing indole-3-butyric acid (IBA) level. Our study indicates that a species-specific variation exists in NPA effect on hypocotyl elongation. Importantly, in addition to influencing PAT, it also has a wider range of influence encompassing modulation of hormonal levels and cellular metabolites.

## Results

### NPA stimulates hypocotyl elongation in tomato seedlings

Light-grown tomato seedlings raised in plastic boxes facilitating air exchange [Lid open boxes (LO)] on NPA containing media displayed elongated hypocotyls. The response was exacerbated in seedlings grown in the boxes with the inverted lid sealed with Parafilm to minimize the air exchange [Lid closed boxes (LC)]. The tomato seedlings grown in LC in light, on varying NPA concentration, displayed stimulation of hypocotyl elongation and reduction in root elongation (Fig. 1A). The epidermal cells of hypocotyls of NPA-treated seedlings were more elongated and less wide than the control seedlings (Fig. 1B). The roots of horizontally placed NPA-treated seedlings were less responsive to gravitational stimulus. Consistent with loss of gravitational sensing, the tips of NPA-treated roots displayed a reduction of amyloplasts staining and were nearly bereft of amyloplasts staining at 1 μM NPA ((Fig. 1C). The possibility that NPA may have affected auxin transport was also indicated by the reduction in the level of DR5::GUS and IAA2::GUS staining in NPA-treated seedlings, signifying reduced activity of auxin in NPA-treated root tips (Fig. 1D). Taken together these results indicated that while NPA affected auxin-mediated responses, it also affected hypocotyl elongation. Considering that NPA-induced hypocotyl elongation was reminiscent of ethylene-mediated hypocotyl elongation, we hypothesized that above response probably resulted from stimulation of ethylene synthesis by NPA.

**Figure 1:**
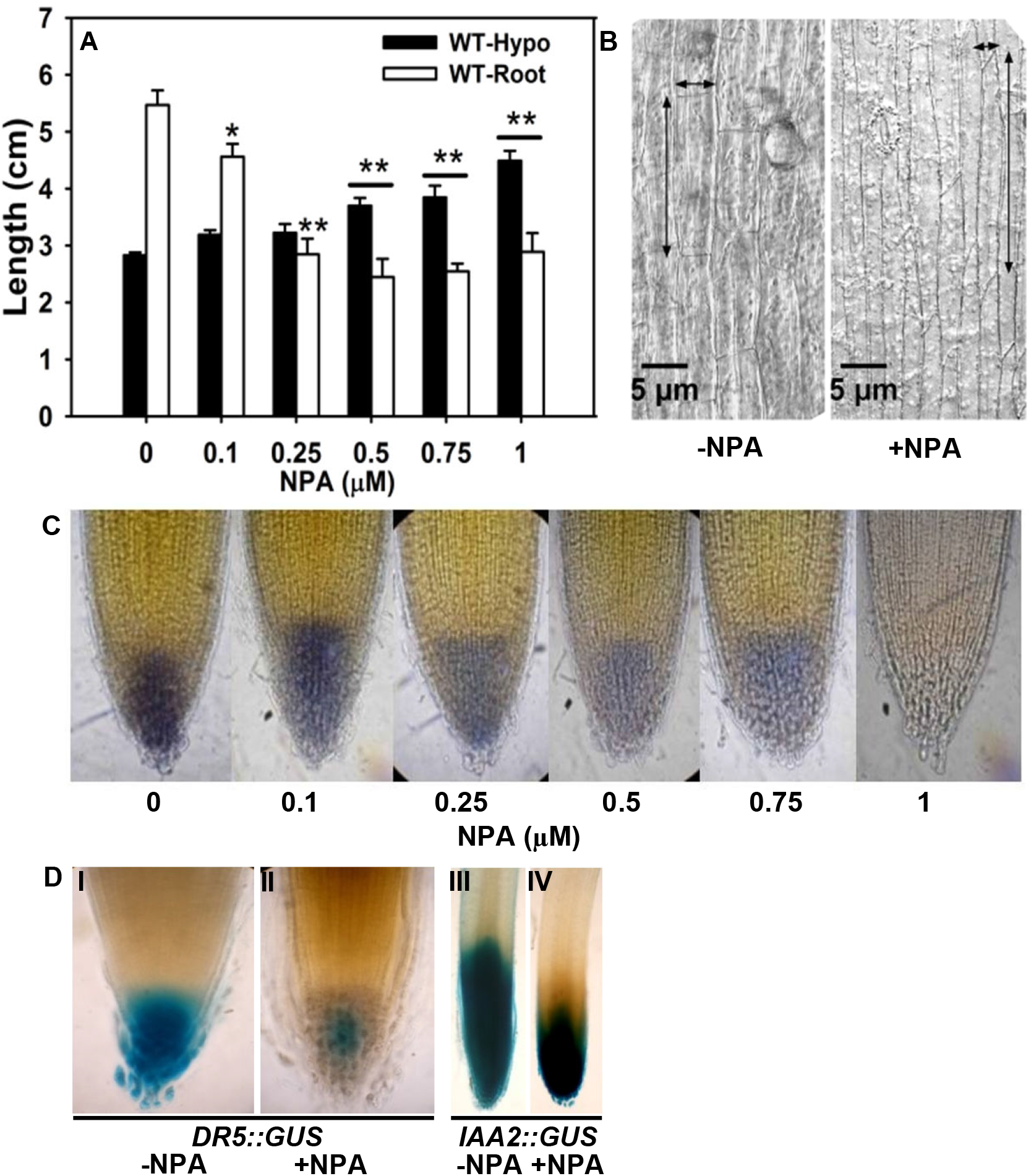
Effect of NPA on hypocotyl and root elongation in light-grown seedlings. **A**- Increasing NPA concentration stimulates hypocotyl elongation and inhibits root elongation. **B**- NPA-treated (1 μM) hypocotyls display elongated and narrow epidermal cells compared to control. **C**- Progressive reduction in amyloplasts levels in root tips with increasing dosage of NPA. **D**- Reduced expression of auxin activity reporters *DR5::GUS* (**I, II**) and *IAA2::GUS* (**III, IV**) in NPA-treated root tips. (n>30 per replicate, ** P < 0.001).

### 1-Methylcyclopropene reverses NPA-stimulated hypocotyl elongation

To ascertain that NPA stimulated ethylene synthesis, the seedlings were grown in LC to confine the endogenously produced ethylene within the box. As a control, NPA-treated seedlings were also grown in LO to permit the escape of ethylene. Interestingly, NPA-stimulated the hypocotyl elongation in both LO and LC, albeit with a lesser magnitude in LO (Fig. 2A). The treatment of seedlings with exogenous ethylene phenocopied NPA-like response with elongated hypocotyl (Fig. 2A) and reduction in root length (Fig. 2B), albeit with lesser magnitude. In contrast, seedlings treated with 5 μM synthetic auxin 2,4-dichlorophenoxyacetic acid (2,4-D), reduced both hypocotyl and root elongation. It is known that mercuric perchlorate (MP) is an efficient absorbent of ethylene and reduces ethylene concentration in closed boxes (Fugate et al., 2010). NPA-treated seedlings grown in presence of MP showed hypocotyl and root length similar to LO. Conversely, 1-methylcyclopropene (MCP), an inhibitor of ethylene action, inhibited hypocotyl elongation and stimulated root elongation, both in NPA-treated and control seedlings. Taken together, these results indicated that the NPA-treatment stimulated ethylene biosynthesis in the tomato seedlings.

**Figure 2:**
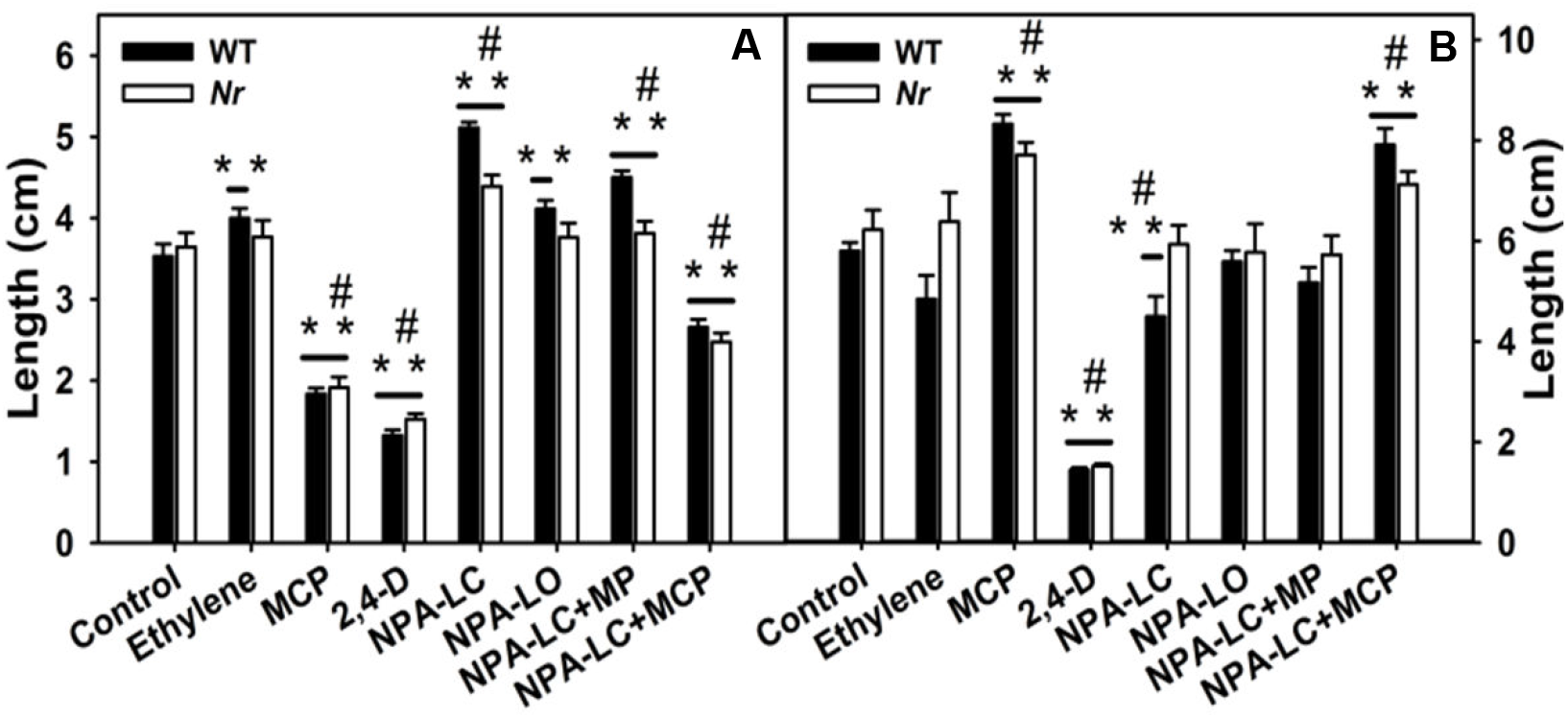
MCP reverses the NPA-stimulated hypocotyl elongation. **A-B**, The effect of ethylene, MCP, 2-4D, and MP alone or along with NPA on hypocotyl (A) and root (B) elongation of tomato or its *Nr* mutant. Concurrent MCP and NPA, or MP and NPA, application partially reverses NPA-effect both in hypocotyl (**A**) and in root (**B**). Asterisk indicates statistically significant difference from control seedlings, and # between ethylene- and NPA-treated seedlings (n = 30 per replicate; Student’s t-test, **/# P < 0.001).

To further ascertain the role of ethylene, the NPA-response was examined in Never-ripe (*Nr*), an ethylene insensitive mutant. Interestingly, NPA-stimulated the hypocotyl elongation in *Nr* mutant too, albeit the response was less than WT (Fig. 2A). It is plausible that *Nr* mutant that has a truncated ETR3, a tomato ethylene receptor (Wilkinson et al., 1995), may be sensing NPA-induced ethylene using other ethylene receptors. Though *Nr* and WT seedlings showed nearly similar hypocotyl elongation with NPA and other treatments (Fig. 2A, B), their effect on *Nr* root was more subdued than WT. Nonetheless, MCP-treatment inhibited NPA-mediated hypocotyl elongation and stimulated root elongation in *Nr* mutant further indicating a role of ethylene.

### NPA stimulates ethylene formation in tomato seedlings

The phenocopying of NPA-treatment with ethylene and its reversal by MCP indicated that NPA stimulated ethylene emission from the seedlings. Consistent with above view, ethylene emission from NPA-treated seedlings was higher than the untreated seedlings (Fig. 3A). MCP-treatment did not affect ethylene emission, as NPA-treated seedlings grown with or without MCP, emitted a similar amount of ethylene. The reduced level of ethylene emission in NPA-treated seedlings grown in LO was due to diffusion of ethylene from the box. The ethylene released from the seedlings was also measured by trapping it with MP and later determining the released ethylene by gas chromatography. In summary, these experiments indicated that NPA stimulated ethylene emission from tomato seedlings (Fig. 3A).

**Figure 3:**
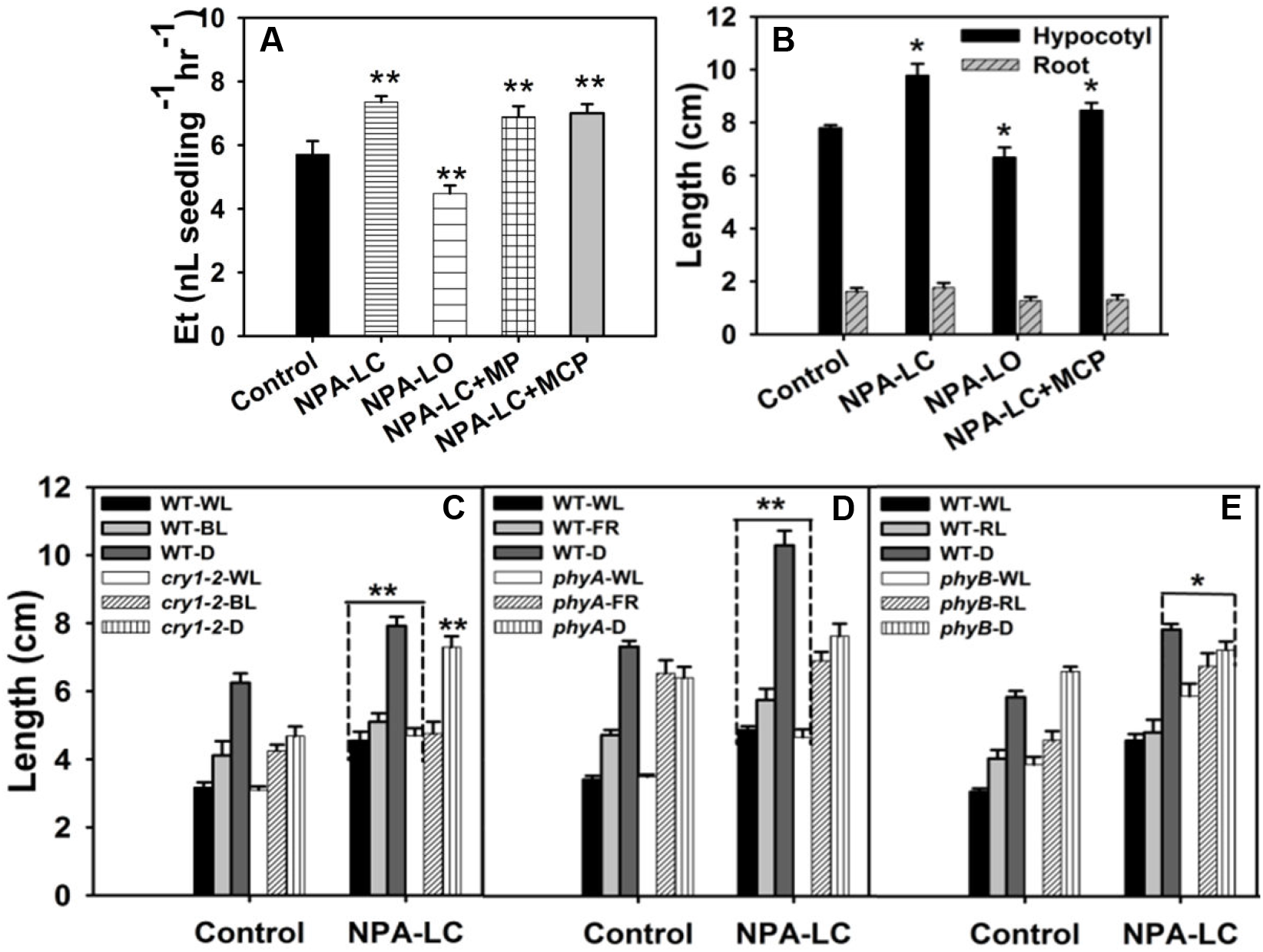
Effect of NPA on ethylene emission; on dark-grown seedling; and photoreceptor mutants. **A**- Ethylene emission from light-grown seedlings with or without NPA. The inclusion of MP or MCP does not affect NPA-induced ethylene emission. **B**- In dark-grown seedlings, NPA affects only hypocotyl elongation. Note MCP partly reverses its action. **C-E**, Photoreceptor mutants *cry1-2* (**C**), *phyA* (**D**), and *phyB1* (**E**) retain NPA-stimulated elongation in white light and darkness. WL= White light, RL= Red light, B= Blue light, D= Darkness. Asterisk indicates statistically significant differences between control and NPA treated seedlings; (n ≥ 15 per replicate; one-way ANOVA; Student’s t-test, * P < 0.05 and ** P < 0.001).

### NPA stimulated hypocotyl elongation even in etiolated seedlings

Several studies have indicated that the ambient light influences hypocotyl elongation in a species-specific fashion (Pierik et al., 2006; Smalle and Van Der Straeten, 1997). In Arabidopsis seedlings, the NPA-mediated inhibition of the hypocotyl elongation is restricted only to light-grown seedlings (Jansen et al., 1998). In contrast to Arabidopsis, NPA-treatment to dark-grown seedlings stimulated hypocotyl elongation in a fashion similar to light-grown seedlings in LC. Analogous to the light-grown seedlings, the inclusion of MCP in LC partially reversed this inhibition (Fig 3B). However, NPA-treated seedlings in LO did not show a stimulation of hypocotyl elongation. Unlike light-grown seedlings, no NPA-mediated inhibition of root elongation was observed in etiolated seedlings.

### Photoreceptor mutants retain NPA-stimulated elongation in white light

It is well established that light inhibits the hypocotyl elongation using multiple photoreceptors; consequently, the loss of individual photoreceptors promotes hypocotyl elongation (Jensen et al., 1998). Therefore, we examined whether tomato cryptochrome1 mutant (*cry1-2*), phytochromeB1 mutant (*phyB1*) and phytochromeA mutant (*phyA*) which are insensitive to blue, red and far-red light respectively, displays NPA-stimulated hypocotyl elongation (Fig. 3C-E). In white light barring *phyB1* mutant that had longer hypocotyl (Fig. 3E), the *cry1-2* and *phyA* mutant hypocotyls were of similar length as of wild-type (Fig. 3C-D). All three mutants displayed NPA-stimulated hypocotyl elongation similar to wild-type control under white light. Consistent with the requirement of full WL spectrum for suppression of hypocotyl elongation, wild-type grown under the blue, red and far-red light, showed elongated hypocotyls than under WL. The NPA-treatment to WT seedlings only marginally stimulated hypocotyl elongation under the blue, red, and far-red light, while the *cry1-2* mutant under blue light and *phyA* mutant under far-red light did not display NPA-stimulated hypocotyl elongation. Interestingly, the NPA-treatment significantly stimulated hypocotyl elongation in red-light grown *phyB1* seedlings. Unlike WT seedlings, NPA did not stimulate hypocotyl elongation in dark-grown *phyA* and *phyB1* mutant, whereas, it stimulated elongation in the *cry1-2* mutant (Fig. 3C).

### NPA-treatment reduces the indole-3-butyric acid level in shoots

Considering that NPA stimulated ethylene emission from tomato seedlings, we next examined whether it influenced the levels of other phytohormones too. Since in light-grown seedlings auxin is transported from cotyledon to hypocotyls (Zheng et al., 2016), we examined the hormonal profiles in whole shoots including cotyledons rather than in the hypocotyls (Fig. 4, Supplemental Table S1). The levels of gibberellic acid and brassinosteroids in shoots were below the limit of detection. The hormonal profiling revealed that NPA did not significantly alter the levels of IAA, methyl jasmonate, abscisic acid, and jasmonic acid in seedlings grown either in LC or LO compared to control seedlings. The sole exception was salicylic acid level, which declined in NPA-LO and zeatin whose level increased in NPA-LO and NPA-LC. In contrast, the NPA-treatment drastically reduced the levels of IBA in seedlings grown both in LC and LO.

**Figure 4:**
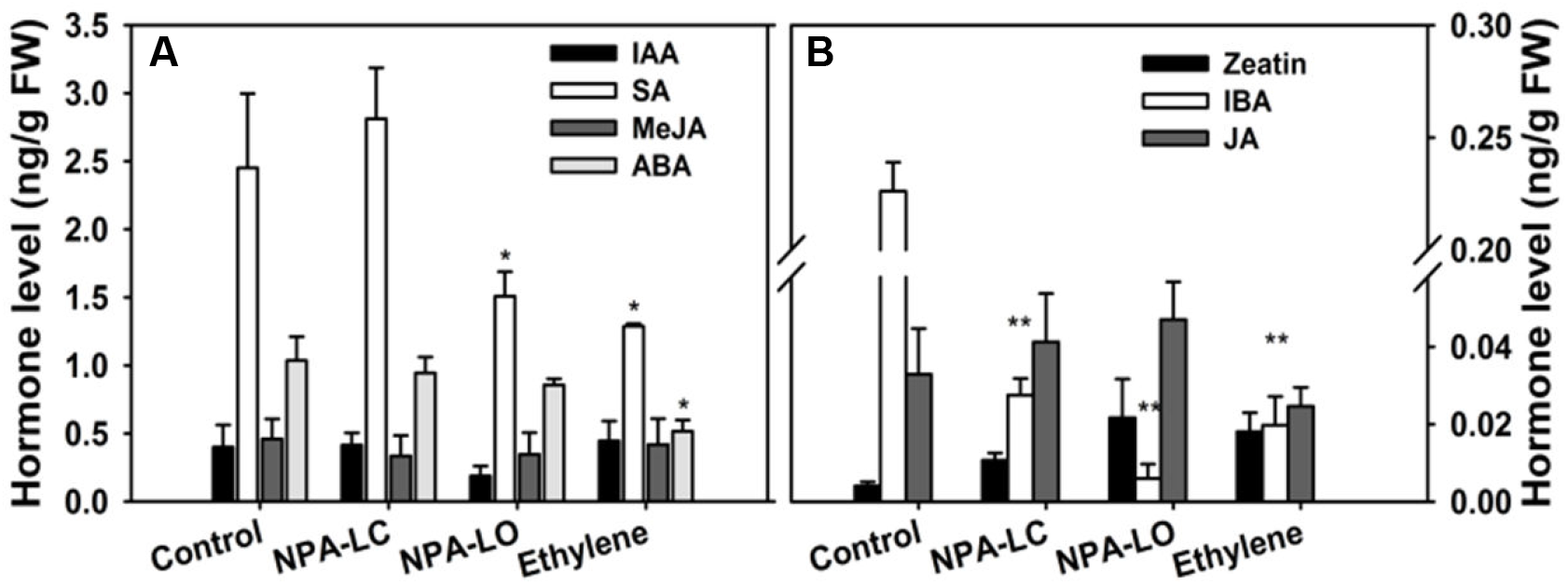
Effect of NPA on level of different phytohormones in shoot of in shoots of NPA-LC-, NPA-LO-, Ethylene-treated and untreated control seedlings. SA-Salicyclic acid, MeJa-Methyl jasmonate, JA- Jasmonic acid, IBA-Indole-3-butyric acid, IAA-Indole-3-acetic acid, ABA-Abscisic acid. The values represent the mean values ± SE of five independent biological replicates (n≥15 per set). one-way ANOVA; Student’s t-test, * P < 0.05 and ** P < 0.001).

### NPA and ethylene affect metabolite profiles in distinctly different fashions

The growth and developmental responses are intimately related to cellular metabolism, which in turn is modulated by endogenous changes of hormones and other regulatory processes. Considering both NPA and ethylene stimulated hypocotyl elongation, we compared the profiles of primary metabolites of NPA- and ethylene-treated seedlings with untreated control seedlings. The GC-MS analysis of the methanolic extract of shoots of 5-days old light-grown seedlings revealed ca. 108 metabolites including isomers (Supplemental Table S2). Among these, the molecular identity could be ascertained only for 103 metabolites. The principal component analysis (PCA) revealed metabolite changes in the seedlings treated with exogenous ethylene were distinctly different from the control seedlings (Fig. 5; Supplemental Table S3). In contrast, the metabolite profiles of NPA-treated seedlings were though distinct, were closer to the control seedlings. It is apparent that though both NPA and ethylene stimulated hypocotyl elongation, they influenced the metabolic profile in very different fashions.

**Figure 5:**
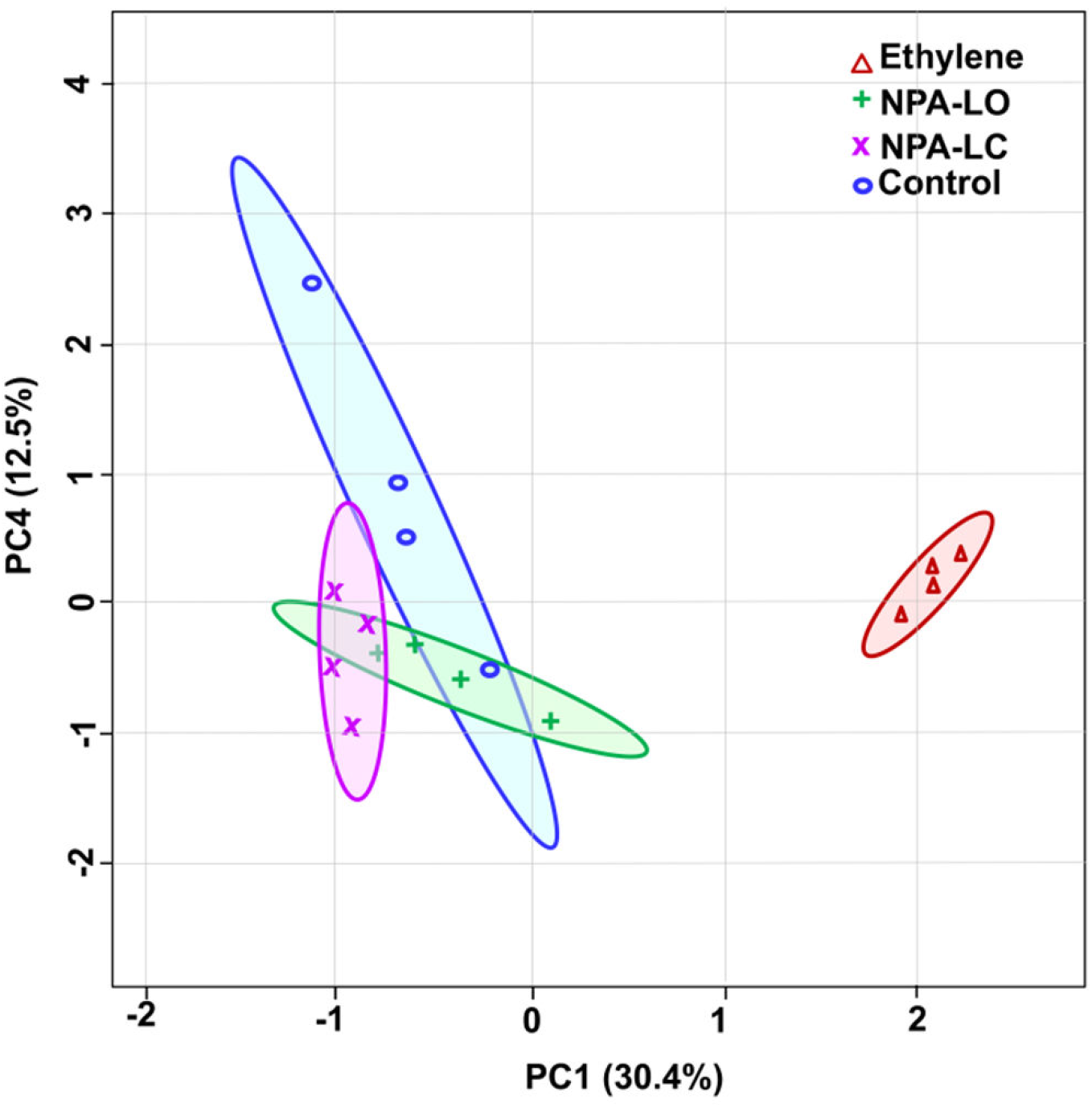
Principal Component Analysis (PCA) of primary metabolite profiles of shoots of 5-day-old light-grown seedlings grown with or without NPA, or with ethylene. The metabolite profile of ethylene-treated seedlings is distinctly different from NPA-treated and control seedlings. The PCA was constructed using MetaboAnalyst 3.0 and the variance of PC1 and PC4 is given within parentheses. Seedlings were grown without NPA (Control), or with NPA (NPA-LC, and NPA-LO), and with ethylene (Ethylene).

The comparison of the levels of individual metabolites from shoots of treated seedlings with control seedlings corroborated that NPA and ethylene actions were distinctly different (Supplemental Figure S1, Supplemental Table S4). Among the 103 detected metabolites, the significantly differential influence of NPA-LC, NPA-LO, and ethylene was discernible for 35, 37, and 54 metabolites respectively (Supplemental Table S5). The exposure to ethylene elevated the levels of citrate, malate, and fumarate in the TCA cycle and lowered the level of isocitrate and succinate (Fig. 6). In contrast, NPA-LC treatment affected these metabolites in subdued fashion. A similar diversity was also manifested in the levels of the majority of amino acids. Exposure to ethylene elevated the levels of glycine, l-leucine, proline, valine, phenylalanine, methionine, cystathionine, asparagine, and aspartate, while treatment with NPA lowered the levels of leucine, lysine, tyrosine, cystathione levels. Consistent with PCA, the NPA-LO influence on the levels of above metabolites was closer to NPA-LC than ethylene. The influence of ethylene on other metabolites like sugars, fatty acids, amines, and other organic acids was also distinct from NPA treatment.

**Figure 6:**
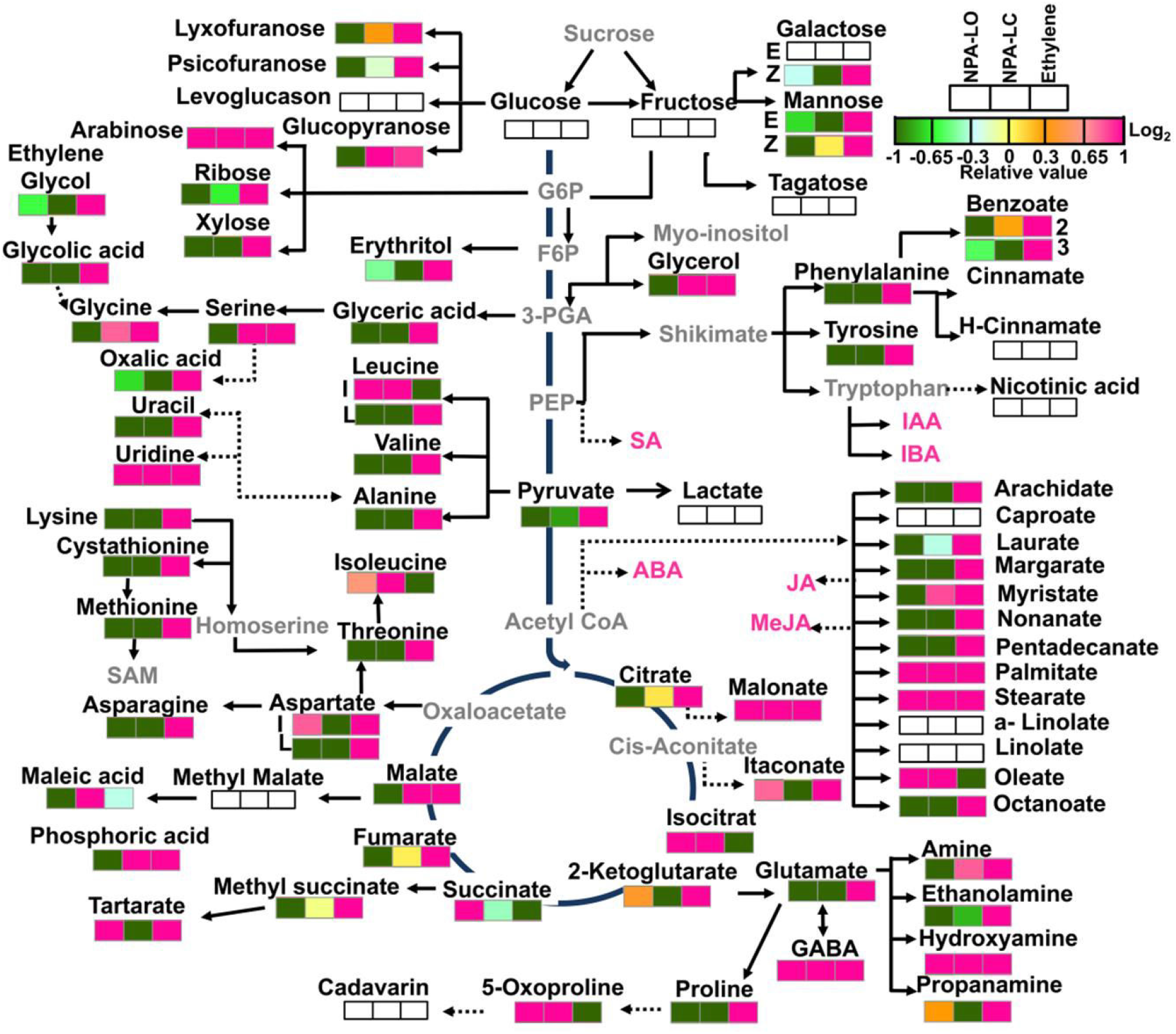
An overview of the metabolic pathways representing relative abundance of metabolites and hormones in shoots of NPA-LC, NPA-LO, Ethylene-treated seedlings compared to untreated control. The abundance of each metabolite and hormone is represented by a heat map. The relative changes in the metabolite levels at ethylene, NPA-LC, and NPA-LO treatments were determined by calculating the treatment/control ratio. Log_2_ -fold changes are represented by varying shades of colors in the boxes with magenta and dark green representing a maximum increase and decrease respectively at upper right-hand corner. The metabolites whose levels were higher or lower to Log2 fold changes (−1 or+1) and statistically significant (Supplemental Table S3) in the treated sample versus non-treated in comparison to control are only shown on the pathway. The white box represents the metabolites, which were detected but not statistically significant. The metabolites in gray letters on pathway were not detected in the GC-MS analysis. The pink lettered metabolites denote hormones measured by LC-MS. The values are the mean ± SE (n≥4).

### Correlation networks are distinctly different in NPA- and ethylene-treated shoots

To decipher the shift in the interaction among different metabolites and with the plant hormones, the correlation networks (r>0.95) were plotted after subtracting interactions that were present in the control seedling (Fig. 7, Supplemental Table S6). The networks were populated with only those metabolites that significantly differed from control seedlings by at least 1.5 fold higher or lower expression. Consistent with a unique PCA profile that significantly differed from control seedlings, the ethylene-treated shoots showed a maximal number of interactions (Network density-0.534, edges 778, nodes 55). Though NPA-LC and NPA-LO differed in their interactions, the treated shoots showed considerably lower unique interactions than the control seedlings (NPA-LC Network density-0.285, edges 94, nodes 35; NPA-LO Network density-0.209, edges 171, nodes 41). Interestingly, the interaction network in ethylene-treated shoots was exclusively populated by positive interactions, whereas in NPA-LC and NPA-LO treatments both positive and negative interactions were present.

**Figure 7:**
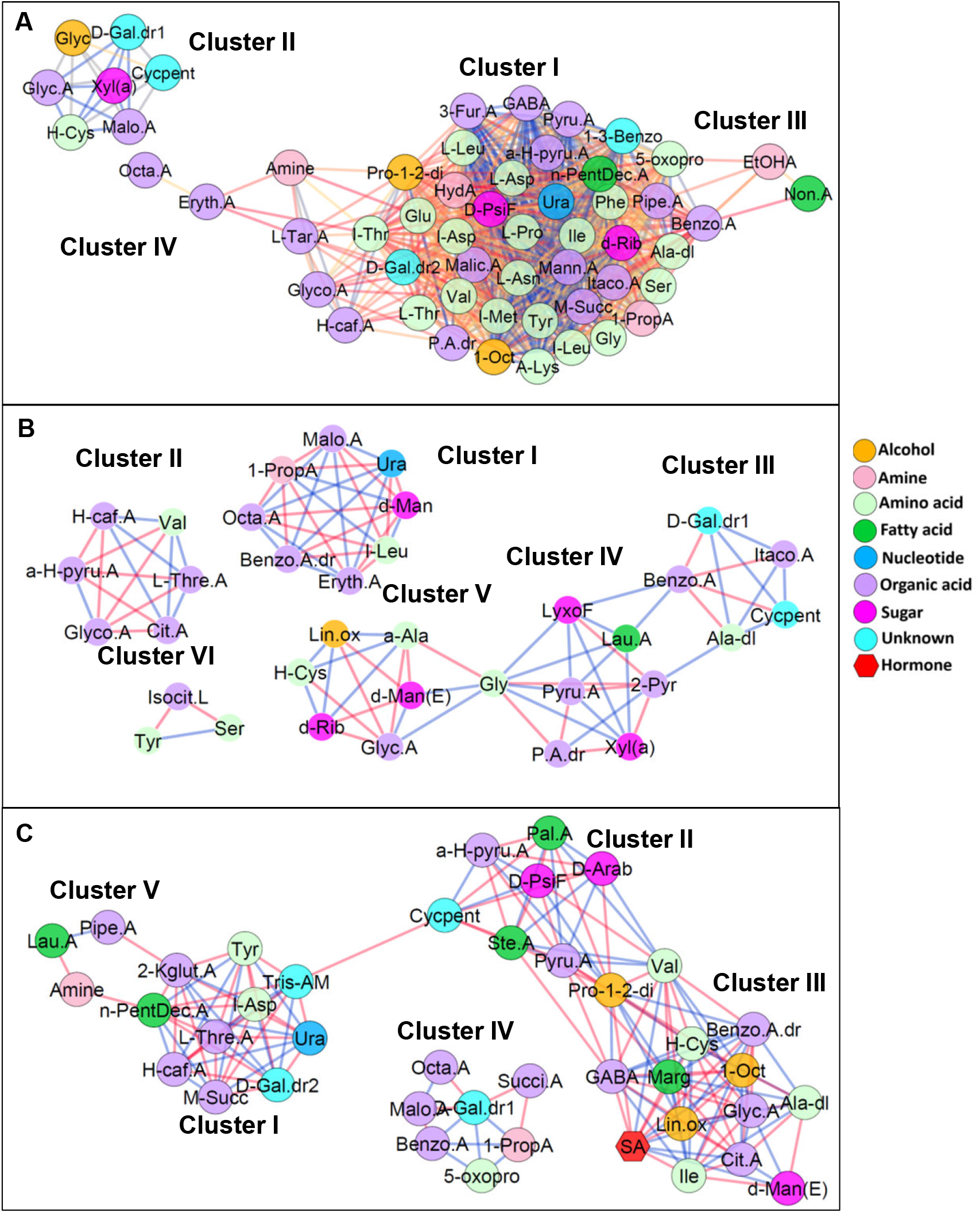
Correlation network of metabolites and hormones in shoots of NPA-LC, NPA-LO, Ethylene-treated seedlings compared to untreated control. The networks were constructed by subtracting control from Ethylene (A), NPA-LC (B), and NPA-LO (C) and including only significantly varying (log 2-fold ≤ −1,>+ 1) metabolites and hormones. Only interactions (p ≤0.05) with r ≥ ±0.95 were used for generating the network. The nodes are represented by different colored circles for metabolites and red colored hexagons for hormone(s). The edges are represented with blue lines for positive correlations and red lines for negative correlations. Statistically significant differences in the level of metabolites under different treatment conditions were determined through One-Way ANOVA (P< 0.05). The full names of the metabolites depicted on the network are given in Supplemental Table S1.

There was little similarity in metabolite interaction between ethylene- and NPA-treated seedlings. Between ethylene and NPA-LC, only 7 interactions were common, while NPA-LO had 18 common interactions with ethylene. In contrast, only 5 interactions were common between NPA-LC and NPA-LO, of these 3 were positive and two were negative interactions. The interaction networks also corroborated the stimulating influence of ethylene on metabolite levels, with all of the examined metabolites showing positive interactions. The interaction profiles also were in conformity with PCA profiles, wherein ethylene-treated shoots made a distinct cluster. While NPA-LO and NPA-LC were closer to untreated seedlings, the difference in their interaction pattern is consistent with their PCA profiles being on either side of untreated seedlings.

Examination of subtraction network between ethylene-treated and control seedlings revealed the densest networks, the network being populated by most amino acids and other metabolites constituting a major cluster. Surprisingly none of the plant hormones that showed variation on ethylene treatment mapped on the network. Majority of the metabolites showed >30 interactions, with malic acid with highest 41 interactions, consistent with it being the part of TCA cycle. Contrastingly, the subtraction network between NPA-LC versus control was sparsely populated and consisted of six clusters, three of which were interlinked whereas three were independent clusters. Contrary to expectation, the NPA-LO versus control subtraction network differed from the NPA-LC versus control. There were only 5 interactions common between these two networks, whereas two interactions flipped, one as positive and other as negative interaction. Interestingly, only in NPA-LO a plant hormone SA was mapped on the network, having a negative interaction with GABA and positive interaction with citric acid.

## Discussion

NPA is one of the widely used phytotropins in the higher plants (Teale and Palme, 2018). The observation that NPA stimulated hypocotyl elongation in a dose-dependent fashion indicated that NPA affected an endogenous process regulating hypocotyl elongation in tomato. The increased elongation likely stems from cell expansion, as the epidermal cells of NPA-treated hypocotyls, were longer but less wide than the control. The observed stimulation of hypocotyl elongation by NPA in tomato is contrary to Arabidopsis, where NPA-treated seedlings grown in white light show reduced hypocotyl length (Jensen et al., 1998). However, this converse NPA effect is confined to hypocotyl elongation, as the NPA-treated tomato seedlings show reduced root elongation similar to Arabidopsis. In consonance with reduced root elongation, root tips of NPA-treated tomato seedlings displayed a reduction in amyloplasts levels, reduced expression of auxin activity reporters; *DR5:GUS* and *IAA2:GUS*, indicating reduced auxin transport.

Ostensibly though NPA inhibited auxin transport in tomato seedlings, it did not elicit a hypocotyl inhibition as observed for Arabidopsis. The possibility that tomato hypocotyls may be insensitive to NPA seems unlikely. The NPA-treatment of tomato apex completely stopped auxin-induced leaf formation indicating that tomato shoots are sensitive to NPA (Reinhardt et al., 2000). An alternative possibility is NPA-inhibition of auxin transport may lead to localized auxin accumulation thus eliciting hypocotyl elongation. This possibility is also unlikely as the application of 2-4D, a synthetic auxin analog, rather than phenocopying the hypocotyl elongation response, lead to a severe reduction of both hypocotyl and primary root length. Moreover, in the light-grown seedlings, auxin mediating hypocotyl elongation is mainly transported from the cotyledons, at least in Arabidopsis, and this transport is sensitive to NPA (Tao et al., 2008). Seemingly, the NPA-stimulated hypocotyl elongation is not related to alteration of auxin-mediated responses.

The hypocotyl elongation in light-grown seedlings is also influenced by ethylene, which conversely to auxin, stimulates the hypocotyl elongation (Smalle et al., 1997). It is plausible that NPA-treatment by blocking auxin transport altered cellular homeostasis, which in turn stimulated ethylene biosynthesis owing to physiological stress. Our results indicate that NPA-induced hypocotyl elongation likely results from its effect on ethylene synthesis. The scrubbing of ethylene by the inclusion of MP reduced the magnitude of hypocotyl elongation in NPA-treated seedlings. Likewise, MCP, a noncompetitive inhibitor of ethylene receptors (Sisler, 2006), that affects hypocotyl elongation in tomato seedlings (Santisree et al., 2011), significantly inhibited the hypocotyl elongation both in NPA-treated as well as in control seedlings. The inhibition of hypocotyl elongation in MCP-treated control seedlings indicate that endogenously produced ethylene plays an important role in hypocotyl elongation in light-grown tomato seedlings. However, the reduction of hypocotyl elongation by MCP was less severe in NPA-treated seedlings most likely due to NPA-stimulation of ethylene synthesis. The exposure of the seedlings to ethylene elicited hypocotyl elongation response similar to NPA albeit with reduced magnitude. The higher magnitude of hypocotyl elongation in NPA-LC than in NPA-LO is likely due to the retention of ethylene in NPA-LC. Consistent with above results, the NPA-treated seedlings emitted a higher level of ethylene and inclusion of MP or MCP did not affect significantly affect ethylene emission from seedlings. Since the response of *Nr* mutant of tomato, which is defective in ETR3 like receptor (Wilkinson et al., 1995), was closely similar to NPA-treated seedlings, it can be assumed that an ethylene receptor other than ETR3 may be involved in above responses.

In Arabidopsis, NPA-inhibition of hypocotyl elongation is observed under white, blue, red, and far-red light (Jansen et al., 1998), but not in dark-grown seedlings. Contrary to this, NPA-treatment elicited hypocotyl elongation in dark-grown tomato seedlings and inclusion of MCP reduced the elongation response. Using photoreceptor mutants, it was suggested that the light-dependent NPA inhibition of hypocotyl elongation in Arabidopsis involves both phytochrome and cryptochrome (Jansen et al., 1998). Unlike Arabidopsis, NPA-treated tomato seedlings grown under blue, red, and far-red lights rather than showing hypocotyl inhibition showed milder stimulation of hypocotyl elongation. Likewise, tomato *cry1-2, phyA*, and *phyB1* mutants retained NPA-induced hypocotyl elongation in white light and darkness, excepting *cry1-2* in blue light, *phyA* in far-red light and *phyB1* in darkness. Considering that NPA stimulated hypocotyl elongation in darkness, and under different lights it either stimulated mild elongation or had no effect in wild-type and photoreceptor mutants; it can be assumed that NPA action in tomato is not related to photoreceptor-mediated modulation of auxin transport (Jansen et al., 1998; Kraepiel et al., 2001; Liu et al., 2011).

Though NPA is widely used as phytotropin, only limited information is available about its mode of action. There is also no prior information whether NPA treatment affects levels of other hormones and/or cellular metabolome. NPA reportedly binds *in vivo* to at least five macromolecules viz., ABCB1, ABCB19, TWISTED DWARF1 (TWD1), aminopeptidase M1 (APM1) and ABP1 (Zhu and Geisler, 2015; Teale and Palme, 2018) and also affects actin polymerization via TWD1 (Zhu et al., 2016). Given the diversity of its putative binding sites and range of the phenotypes elicited, it is likely that NPA displays its effect by action on several cellular responses. The NPA-elicited hypocotyl elongation is a cumulative manifestation of a cascade of cellular/metabolic responses, which also involves the participation of plant hormones. Considering that NPA stimulated ethylene emission from the seedlings, it is plausible that it influenced the levels of other hormones too. The higher level of zeatin in NPA-treated hypocotyls corroborates that NPA had wide-ranging effects. In Arabidopsis, zeatin stimulates hypocotyl elongation in light (Saibo et al., 2007) therefore; it may have an analogous role in tomato.

Surprisingly, NPA did not influence IAA level but drastically reduced IBA level. In Arabidopsis, NPA does not affect IBA basipetal/acropetal transport in the root and basipetal transport in the hypocotyl (Rashotte et al., 2003), presumably IBA transport in tomato is similarly insensitive to NPA. Considering that the IAA level was not altered, it is unlikely that the reduced IBA level was due to enhanced transport of IBA. It is more likely that the reduction of IBA level arose from its conversion to IAA (Bartel et al., 2001; Strader and Bartel, 2011). Nonetheless, the reduced IBA level appears to be causally related to NPA-induced hypocotyl elongation (Ludwig-Muller, 2000). Emerging evidence indicates that IBA too has a role in growth and development as IBA resistant *rib1* mutant shows alteration in growth, development, and response to exogenous auxin (Poupart et al., 2005). The IBA-derived IAA has also been shown for its importance in organ development (Strader et al., 2010, 2011; Frick and Strader, 2018). There seems to be an interrelationship between reduced IBA levels, enhanced zeatin level, and higher emission of ethylene in NPA-treated hypocotyls. The exposure of seedlings to ethylene also elicited the reduction in IBA and increase in zeatin levels supporting such an interrelationship. While enhanced hypocotyl elongation may involve a collective action of these hormones, the shifts in their levels also imply that NPA modulates cellular homeostasis in tomato.

Consistent with this assumption, NPA elicited a shift in the levels of several metabolites. The PCA profiles of NPA-treated hypocotyls were distinctly different the control, and even NPA-LC and NPA-LO treatments had different PCA profiles. Interestingly, ethylene-treated hypocotyls displayed drastically different PCA profile than NPA-LC and NPA-LO treatments. This difference may stem from the likelihood that endogenously produced ethylene may influence the metabolic responses in a different way than the exogenously applied ethylene. Moreover, it is plausible that the NPA-induced ethylene emission may be restricted to specific tissues/organs, whereas exogenous ethylene most likely will globally affect all tissues/organs. While the information for tissue-specific ethylene synthesis is not available in tomato, in Arabidopsis ACO2 mRNA is predominantly localized in elongating cells of apical hook and its expression is correlated with ethylene-induced hook curvature (Raz and Ecker, 1999). In addition, while few of NPA-mediated metabolite changes may involve ethylene, it may also independently of ethylene influence other metabolites. While these possibilities remain, it is difficult to distinguish between the metabolite changes mediated by NPA via ethylene and/or independently of ethylene. However, the diversity between ethylene and NPA influence on metabolite levels support above assumption, as the shifts seen in the levels of individual metabolites by NPA and ethylene are very different. While ethylene increased the levels of most of the amino acids, NPA lowered the levels of several amino acids. Similar diversity between NPA and ethylene action was also observed for amines, sugars, and fatty acids. Considering that abiotic stress induces accumulation of amino acids such as valine, leucine, threonine, and methionine (Obata and Fernie, 2012), it can be assumed the observed accumulation of amino acids in ethylene-treated seedlings may represent an analogous stress condition.

One of the way to distinguish between overlapping and independent action of NPA and ethylene is to compare the correlation networks (Langfelder and Horvath, 2008). Ostensibly, the differences in their action on metabolite and hormonal levels should be similarly reflected in the correlation networks. Consistent with its distinctly placed PCA profiles, the network of ethylene treated plants were denser, but shared very few interactions with NPA-LC or NPA-LO. The examination of correlation networks revealed that the NPA-LC moderately influenced metabolic homeostasis with reference to control as evident by relatively few unique interactions. Interestingly, the NPA-LO shared only five interactions with NPA-LC and rest were unique. This is also consistent with the proximity of PCA profiles of NPA-LC and NPA-LO with control and at the same time, their positioning on PCA plot is on two different flanks of control. While the NPA treatment is identical, in NPA-LC, the released ethylene is confined to the box, whereas in NPA-LO, it escapes. Moreover NPA-LC also likely to have lesser oxygen due to its consumption in respiration. These two factors may have contributed to these differences in PCA profiles. Nonetheless, NPA-LO shows only slightly lesser hypocotyl elongation than NPA-LC, the contribution of above two processes on elongation seems to be minimal.

An intriguing observation was that none of the detected plant hormones mapped on the correlation networks, barring salicylic acid in NPA-LO. This ambiguity reflects that fact that these correlation networks are not robust as these represent only primary metabolites which numbered <100, whereas, in tomato so far 2677 metabolites have been annotated in TOMATOCYC version 3.5 (https://www.plantcyc.org/databases/tomatocyc/3.5). Hormonal influence on metabolic homeostasis may be more subtle and directed towards specific metabolites influencing growth such as cell wall expansion observed for NPA-treated hypocotyls. Moreover, our analysis represents only a snapshot of the metabolites, while cellular metabolism undergoes dynamic changes during the time course of hypocotyl elongation.

Notwithstanding these caveats, NPA in addition to being phytotropin also influences other cellular processes. It remains to be determined why tomato hypocotyls respond differently to NPA, while its roots respond similarly to Arabidopsis. The circumstantial evidence indicates that NPA-elevated ethylene may have a role in hypocotyl elongation, as MCP application and MP application reduces hypocotyl elongation. The lesser magnitude of hypocotyl elongation in ethylene-treated plants likely reflects that excess ethylene may be perceived as the stress hormone, as evident by the drastic shift in metabolic profiles (Obata and Fernie, 2012). Considering that all interactions elicited by ethylene were positive, it indicates boosting of overall metabolic flux, a characteristic of plants under stress (Obata and Fernie, 2012). The opposing changes of zeatin and IBA may also have significance, as zeatin reportedly promotes hypocotyl elongation (Saibo et al., 2007). Though not yet conclusively proven, it is suggested the IBA may also have an independent role in growth and development (Ludwig-Müller, 2000). In earlier studies, the growth promoting effect of NPA on pea stem sections were ascribed to it being a weak auxin (Keitt and Baker, 1966; Katekar and Geissler, 1980), which in all likelihood may be related to its effect on levels of other hormones.

In summary, NPA also influences other metabolic responses in addition to acting as a phytotropin. Its influence on other plant hormones may arise from its primary action on auxin transport as well as from its independent action on metabolic responses. Considering the difference in its action in tomato and Arabidopsis, a broader species/organ-specific evaluation of NPA action is desired.

## Materials and Methods

### Plant Materials and Growth Conditions

Wild-type tomato cv. Ailsa Craig *(Solanum lycopersicum), Never-ripe* mutant *(Nr)* in Alisa Craig background; *phyA, phyB1*, and *cry1-2*, mutants in Moneymaker background were used in this study. The seeds were surface sterilized with 20% (v/v) sodium hypochlorite for 10 minutes, washed with distilled water, and sown on wet blotting papers in germination boxes. Seeds were germinated in the darkness at 25±2°C. After the emergence of radicle seeds were overlaid on moist coconut peat (Karnataka Explosive Ltd., Bangalore, India) filled in Petri plates (8.5 cm *l* × 8.5 cm *b* × 1.2 cm *h*). The plates were placed inside transparent plastic germination boxes (9.5 cm *l* × 9.5 cm *b* × 5 cm *h*, Laxbro, Pune). Plants were grown at 25±2°C under 16 hour light (100 μmole m^−2^ s^−1^), 8 hour dark cycle for 5 days under lid closed (LC) condition unless otherwise mentioned. All experiments were independently repeated three or more times (n> 3) using 10 or more 5-day-old seedlings.

### Treatment with NPA and other chemicals

The boxes used in the study had a specially designed lid (Supplemental Figure S2). One face of the lid has a 1.2 cm long and 2 mm tall raised strip in the center of all four side of the lid. The raised strip facilitated the free exchange of gases by leaving a gap between the rim of the box and the lid (Lid open boxes-LO). On another face, the lid lacked the raised strip allowing tightly sealing of the boxes. The lids placed in this orientation were further sealed with Parafilm to minimize the air exchange (Lid closed boxes-LC). For treatment involving ethylene injections, in the center of the lid, a circular hole was made and was sealed with a septum. The septum facilitated injection of ethylene and the withdrawal of the head space for determination of ethylene.

For treatments involving N-1-naphthylphthalamic acid (Sigma Aldrich) and other chemical treatment, chemicals at the concentrations specified were added to the coconut peat prior to overlaying of the germinated seeds. For combined treatments of NPA and 5 μM 2,4-dichlorophenoxyacetic acid (2,4-D) (Sigma Aldrich), both solutions were added at the specified concentrations to the coconut peat. For experiment involving 1-methylcyclopropene (MCP), 30 mg of MCP-releasing powder (0.14% of active ingredient by weight, SmartFreshTM, Rohm and Haas, Springhouse, PA, USA) was dissolved in 5 mL of water in a plastic vial to provide a final gas concentration of 2 μL L^−1^ MCP (Tassoni et al., 2006). The vial containing MCP was taped to the side of the germination boxes and the lid was tightly sealed. The exogenous ethylene treatment, ethylene was injected into tightly sealed boxes to provide a final gas concentration of 1 μL L^−1^.

### Determination of Ethylene Evolution

For measuring ethylene evolution, seedlings were grown in boxes with a septa allowing withdrawal of headspace. After 5 days from germination, one mL of the headspace was withdrawn from the boxes through the rubber septum using a syringe. The headspace gas was then injected into a gas chromatograph equipped with Poropak T column and a flame ionization detector (GC-17A, Shimadzu, Kyoto, Japan) to determine ethylene levels. The estimated ethylene was normalized to the number of seedlings and the duration of the incubation.

### Ethylene trapping by mercuric perchlorate

To reduce the ethylene concentration in the boxes, ethylene scrubbing agent mercuric perchlorate (MP) was used. For trapping ethylene, a cotton ball was placed inside an Eppendorf tube, to that 500 μL of 0.25 M MP was added and Eppendorf tube was tapped to the side of the box and box was tightly sealed. After 5 days of germination, the cotton balls soaked with MP were removed and placed inside a gas-tight plastic box with a rubber septum. Ethylene trapped by MP soaked cotton ball was released using 2 ml of 4 N sodium chloride. The boxes were then incubated for 5 minutes with occasional shaking. One ml volume of headspace was removed for determining the amount of ethylene using the protocol described above.

### The light source and light treatment

Different spectral lights were obtained by using blue (λ_max_ 470 nm), red (λ_max_ 660 nm) (Kwality Photonics, Hyderabad, India), and far-red (λ_max_ 730 nm) (Rothner, Vienna, Austria) LEDs respectively. The fluence rates of blue, red, and far-red light were 5 μmole m^−2^ sec^−1^, 5 μmole m^−2^ sec^−1^, and 15 μmole m^−2^ sec^−1^ respectively.

### Hypocotyl, Root, and Epidermal Cell Length Measurement

To determine root and hypocotyl elongation, seedlings were horizontally laid on a plane surface and lengths were measured using a standard scaled ruler. For the measurement of epidermal cell length, the hypocotyls were cleared overnight using 50% (v/v) lactic acid. The hypocotyls were photographed imaged using Olympus BX 51 microscope equipped with Olympus C7070 camera. The hypocotylar cell lengths were measured using ImageJ software (https://imagej.nih.gov/ij/).

### Metabolite profiling

The metabolite profiling was performed essentially following the protocol of Roessner et al., (2000). For metabolite analysis, the shoots of 5 days old seedlings were excised at the junction of hypocotyl to root and were snap frozen in liquid nitrogen. The frozen shoots were homogenized in liquid nitrogen to a fine powder using mortar and pestle and homogenates were stored at −80□C until further analysis. For determination of metabolites, aliquots measuring 100 mg homogenates were used. The processing and sample derivatization was carried out as described by Bodanapu et al., (2016). The derivatized sample was transferred to a GC-MS injection vial and analyzed by Leco-PEGASUS GCXGC-TOF-MS system (Leco Corporation, USA) equipped with 30 m Rxi-5ms column with 0.25 mm i.d. and 0.25 μm film thickness (Restek, USA). The injection temperature, interface, and ion source were set at 230°C, 250°C, and 200°C respectively. For the separation of groups of metabolites, the run program was set as following; isothermal heating at 70°C for 5 minutes, followed by 5°C min–1 oven temperature ramp to 290°C and final heating at 290°C for 5 minutes. The carrier gas (helium gas) flow rate was set to 1.5 mL/minute. A 1 μL of sample was injected in split less mode and mass spectra were recorded at 2 scans/sec within a mass-to-charge ratio, range 50 to 600.

The raw data were processed by ChromaTOF software 2.0 (Leco Corporation, USA) and metabolite identification was carried out with the NIST MS Search v 2.2 software (National Institute of Standards and Technology) GC/MS Metabolomics library software (2011) (NIST, Department of Commerce, USA) and Golm Metabolome Database libraries (http://gmd.mpimp-golm.mpg.de/). The compound hits which showed maximum matching factor (MF) value (>600) and least deviation from the retention index (RI) was used for metabolite identity. The data were analyzed by normalizing with the sample fresh weight and the internal standard ribitol.

### Determination of phytohormones

The phytohormone extraction from the shoots was performed as described in Pan et al., (2010). An aliquot of 100 mg homogenized powder from shoot sample was used for each experiment. The simultaneous hormonal extraction was done as previously described procedure (Bodanapu et al., 2016). The processed and dried residue was dissolved in 70 μL of precooled 100% methanol followed by centrifugation at 13,000*g* for 5 minutes. The sample was transferred to an injection vial and analyzed using UPLC/ESI-MS (Waters, Milford, MA USA).

The system consists of an Aquity UPLC™ System, quaternary pump, and autosampler. For separation of hormones, the sample was analyzed on a Hypercil GOLD C18 (Thermo Scientific) column (2.1 × 75 mm, 2.7 μm). A gradient elution program was performed using two solvents system, solvent A- containing ultrapure water with 0.1% (v/v) formic acid, solvent B-containing acetonitrile with 0.1% (v/v) formic acid and run for 9 min at 20°C. The abscisic acid (ABA), jasmonic acid (JA), and salicylic acid (SA) detection was performed on ExactiveTMPlus Orbitrap mass spectrometer (Thermo Fisher Scientific, USA) in all ion fragmentation (AIF) modes (range of m/z 50-450) equipped with heated electrospray ionization (ESI) in negative ion mode. The zeatin, IAA, IBA, epibrassinosteroids (Epi-BR) and methyl jasmonate (MeJA) were analyzed in positive ion mode. For both modes, the following instruments setting were used, capillary temperature-350°C, sheath gas flow (N2) 35 (arbitrary units), AUX gas flow rate (N2) 10 (arbitrary units), collision gas (N2) 4 (arbitrary units) and the capillary voltage 4.5 kV under ultra-high vacuum 4e-10 mbar. The hormones were analyzed from the at least 4 different biological replicates of each treatment (NPA, Ethylene) and control. The quantification of each hormone was carried out by comparing the peak areas with those obtained for the respective hormone standards.

### Statistical analysis

To visualize the overall clustering of metabolites and hormones, Principal Component Analysis (PCA) was carried out using Metaboanalyst 3.0 (Xia et al., 2015; http://www.metaboanalyst.ca/). At least three to four biological replicates were used and differences in metabolites accumulation were presented in a two-dimensional graphical display.

### Metabolic Pathway Mapping

The metabolites and hormones detected were mapped on manually constructed general metabolic pathways based on the information available from the published literature and network database as described in the KEGG (Kyoto Encyclopedia of Genes and Genomes, http://www.genome.jp/kegg/) and LycoCyc (Sol Genomic networks, http://www.solcyc.solgenomics.net). To compare the levels of each metabolite across different treatments (Control, Ethylene, NPA-LC, and NPA-LO), Student’s t-tests were performed using statistical analysis of all the metabolites and scored on Microsoft Excel sheet (Tsugawa et al., 2015). Relative average fold changes of metabolites occurring during NPA treatment Vs control and NPA treatment vs ethylene (1 μL L^−1^) were shown on a primary metabolite pathway as presented in Bodanapu et al., (2016) and Do et al., (2010). The replicates with a significance P ≤ 0.05 were used to plot network to highlight patterns of changes in all the conditions of treatment.

### Metabolic and Hormonal networks

To visualize a correlation map between metabolite-metabolites and/or metabolites-hormones, correlation networks were constructed using Cytoscape software version 3.4.0 (http://www.cytoscape.org/; Cline et al., 2007). For correlation networks analysis, four-five biological replicates of treatment were taken. Nodes represent the metabolites (circle), hormones (hexagon shape) and edges represent connectivity between the two metabolites. The overall metabolites scored were pairwise correlated using the Pearson’s Correlation coefficient (PCC). The associations with PCC ± ≤ 0.95 value either in the positive or negative mode were used to create the connectivity between two nodes in the network. For a clearer understanding of the interaction among metabolites and hormones in a network, it was divided into the clusters using Cytoscape plugin, clustermaker 2 version 0.9.5. MCODE algorithm was used for making network and clusters. New correlations in the NPA-LC, NPA-LO and ethylene networks, which were not present in Control, were considered as new associations in treatments or vice-versa.

## Supplemental Data

The following supplemental materials are available.

**Supplemental Table S1**. Hormones detected by LC-MS in shoots of control, NPA-treated (LC and LO) and ethylene-treated seedlings.

**Supplemental Table S2**. Metabolites detected by GC-MS in shoots of control, NPA-treated (LC and LO) and ethylene-treated seedlings.

**Supplemental Table S3**. List of metabolites, PC1, and PC4 loading score for principal component analysis.

**Supplemental Table S4**. Relative changes in the levels of metabolites in NPA-LC, NPA-LO, and ethylene-treated seedlings compared to the control seedlings.

**Supplemental Table S5**. Metabolites in NPA-LC, NPA-LO, and ethylene-treated seedlings that significantly differed from the control.

**Supplemental Table S6**. List of metabolites interactions in ethylene/control, NPA-LC/control, and NPA-LO/control correlation networks.

**Supplemental Figure S1**. Heat maps showing relative expression of metabolites in shoots of NPA-LC, NPA-LO, ethylene-treated seedlings compared to untreated control

**Supplemental Figure S2**. The photographs of the plastic box and the lid used for the experiments.

**Author contributions**
S.N., Y.S., and R.S. Conceived and designed the experiments; S.N. Performed the experiments; S.D. carried out Cytoscape analysis; S.N., R.S., Y.S. analyzed the data and wrote the paper.

